# Diverse bacteriohemerythrin genes of *Methylomonas denitrificans* FJG1 provide insight into the survival and activity of methanotrophs in low oxygen ecosystems

**DOI:** 10.1101/2024.01.07.574550

**Authors:** Cerrise Weiblen, K. Dimitri Kits, Manuel Kleiner, Dominic Sauvageau, Lisa Y. Stein

**Affiliations:** Department of Biological Sciences, University of Alberta, Edmonton, AB, Canada; Department of Plant and Microbial Biology, North Carolina State University, Raleigh, NC, USA; Department of Chemical and Materials Engineering, University of Alberta, Edmonton, AB, Canada

## Abstract

Proteobacterial methanotrophs are dependent on the oxidation of methane for ATP production and assimilation of carbon into biomass. Interestingly, some types of gammaproteobacterial methanotrophs thrive in oxygen-depleted zones of lakes and other aquatic ecosystems despite their reliance on oxygen to support methane oxidation. The model gammaproteobacterial methanotroph, *Methylomonas denitrificans* FJG1, oxidizes methane coupled to nitrate reduction under hypoxia and highly upregulates its expression of bacteriohemerythrin (Bhr). Bhr is a homolog of eukaryotic hemerythrin, which is a protein associated with oxygen binding. Ten *bhr* homologs were identified in the genome of *M. denitrificans* FJG1, requiring phylogenetic and gene expression analyses to pinpoint which homolog is likely responsible for delivering oxygen to support methane oxidation to under low oxygen conditions. This study examined the prevalence and phylogeny of the 10 *bhr* homologs from *M. denitrificans* FJG1 in the genomes of other methanotrophs and across the Bacteria. The homolog denoted “*bhr-*00” was specific to methanotroph genomes, was highly expressed in *M. denitrificans* FJG1 under hypoxia, and its predicted structure was nearly identical to a purified oxygen-scavenging hemerythrin protein from *Methylococcus capsulatus* Bath. Other *bhr* homologs upregulated from denitrifying cultures of *M. denitrificans* FJG1 included those with gene neighborhoods related to oxygen sensing, denitrification and chemotaxis. Together, this study uncovered potential multifunctional roles of bacteriohemerythrin genes of *M. denitrificans* FJG1 under low oxygen conditions and identified the *bhr* homolog that most likely enables and supports oxygen delivery to methane monooxygenase enzymes in anoxic ecosystems.

**Importance:** Aerobic gammaproteobacterial methanotrophs can survive and grow in anoxic lakes, but mechanisms that provide them with oxygen to support methane oxidation remain uncharacterized. *Methylomonas denitrificans* FJG1 encodes 10 copies of bacteriohemerthyrin (*bhr*), of which 7 are expressed at the mRNA level under low oxygen conditions. Comparing the 10 *bhr* homologs from *M. denitrificans* FJG1 with those from other methanotrophs and bacterial genomes shows that two are specific to methanotrophs. Gene neighbourhoods surrounding conserved *bhr* genes in methanotrophs suggest a range of potential functions including oxygen respiration, oxygen sensing, chemotaxis, and nitrate reduction. The results from this study illuminate a previously undescribed diversity of structures and potential functions of *bhr* homologs in *M. denitrificans* FJG1 and related methanotrophic bacteria. The results pinpoint a methanotroph-specific homolog, *bhr*-00, that is likely responsible for oxygen binding and delivery to methane monooxygenase enzymes to promote methane oxidation in low oxygen ecosystems.

## Introduction

Hemerythrin (Hr), and its bacterial homolog bacteriohemerythrin (Bhr), is a multifunctional and structurally diverse protein family linked to multiple metabolic functions in eukaryotes and prokaryotes, including reactive oxygen and nitrogen species detoxification, metal detoxification, iron storage, transmembrane signaling, oxygen sensing, chemotaxis, biofilm formation, cellular respiration, and oxygen binding and transport (1, 2). Previous research has shown that genetically similar Hr and Bhr enzymes perform an array of different functions, placing Hr and Bhr proteins at the crossroads of multiple pathways, even within a single organism. For instance, Baert et al. (3) identified Hr from the leech *Theromyzon tessulatum* which was classified as an iron storage molecule named “ovohemerythrin.” In bacterial systems, Kendall et al. (4) investigated the functions of three Bhr homologs in *Campylobacter jejuni*: one had a protective role in preventing molecules with iron-sulfur clusters (Fe-S) from incurring oxygen damage and the other two were related to flagellar biosynthesis and expression of sigma-factor 28. In the bacterium *Neanthes diversicolour*, a Bhr was found associated with cadmium detoxification functions (5). Research from Xiong et al. (6) showed that the oxygen-sensitive bacterium *Desulfovibrio vulgaris* used a Bhr-like protein as an oxygen sensor with a role in chemotaxis. French et al. (7) showed that Bhr in *Vibrio cholerae* was part of the cyclic-di-GMP signaling system with involvement in biofilm formation.

Several reports of Bhr proteins in gammaproteobacterial methanotrophs have emerged in recent years. These aerobic bacteria use methane as a sole energy and carbon source. A previous survey of *bhr* homologs in methanotroph genomes showed that 80% of those analyzed possessed at least one, and some up to three, *bhr* gene copies (8). In the model strain *Methylococcus capsulatus* Bath, *in vitro* studies showed that purified Bhr could supply oxygen to membrane fractions enriched with particulate methane monooxygenase (pMMO) enzymes, and a crystal structure of the Bhr protein was resolved (9–11). In *Methylotuvimicrobium alcaliphilum* 20Z, Nariya and Kalyuzhnaya (12) showed that overexpression of Bhr supported aerobic respiration but did not enhance methane oxidation capacity under hypoxia or anoxia. A *bhr* knock-out strain of *Methylomonas* sp. LW13 showed decreased growth in a hydrogel with a methane-oxygen counter gradient, but showed no growth impairment in planktonic cultures (13). These studies indicate a relationship between Bhr and oxygen binding, delivery, and/or respiration in methanotrophic bacteria, but none showed the range of potential Bhr activities or their direct roles in supporting methanotrophic metabolism under low oxygen.

Aerobic methanotrophs can use alternate terminal electron acceptors including nitrate (14), nitrite (15), and iron oxides (16). They can also link methane oxidation to fermentation pathways in the absence of oxygen (17). However, there is no known substitute for oxygen that can facilitate the turnover of methane by pMMO enzymes. The present study examines the phylogeny, gene neighborhoods, expression levels, and predicted structures of the 10 *bhr* homologs identified in the genome of *M. denitrificans* FJG1, a denitrifying gammaproteobacterial methanotroph (14), as it transitions from oxic to hypoxic growth conditions. The results highlight a wide diversity of *bhr* contexts in *M. denitrificans* FJG1 and other methanotroph genomes and pinpoints the *bhr* homolog that likely enables gammaproteobacterial methanotrophs to access oxygen to thrive in low oxygen environments.

## Results

### Phylogeny of *bhr* genes identified in *Methylomonas denitrificans* FJG1

*M. denitrificans* FJG1 encodes 10 *bhr* homologs annotated as “hemerythrin family,” “hemerythrin domain-containing protein,” or “bacteriohemerythrin” in its genome (Table 1). The genes, numbered *bhr-00* through *bhr-99*, were designated based on their proximity to the origin of replication in the *M. denitrificans* FJG1 genome sequence (Supp. Fig. S1). Eight of the 10 *bhr* genes are approximately 400 bp, typical of the oxygen-carrying Hr and Bhr proteins originally discovered in eukaryotes (2), whereas *bhr-66* is approximately 1,000 bp and *bhr-55* is approximately 4,000 bp (Table 1). BLASTn searches were performed with each of the 10 *bhr* genes to identify homologs across methanotrophic and non-methanotrophic bacterial genomes. Candidate genes from these BLASTn searches were aligned to a concatenated sequence of the 10 *bhr* homologs (separated by 100 bp spacers to prevent mismatching) from *M. denitrificans* FJG1. and a phylogenetic tree was generated (Fig. 1). The resulting unrooted phylogram of positive matches revealed that *bhr-66* is the most common *bhr*-containing gene found among a diverse assemblage of Bacteria (Fig. 1). In contrast, *bhr-00* and *bhr-99* appear nearly exclusive to methanotrophs in the Methylococcales order. Closer examination of complete *bhr* genes from methanotroph genomes revealed that *bhr-*00 and *bhr*-66 homologs were the most common, but only *M. denitrificans* FJG1 encoded *bhr-*11*,-*33,-44 and -55 among the proteobacterial methanotrophs (Fig. 1; Supp. Data, Sheet 4).

**Figure 1.**
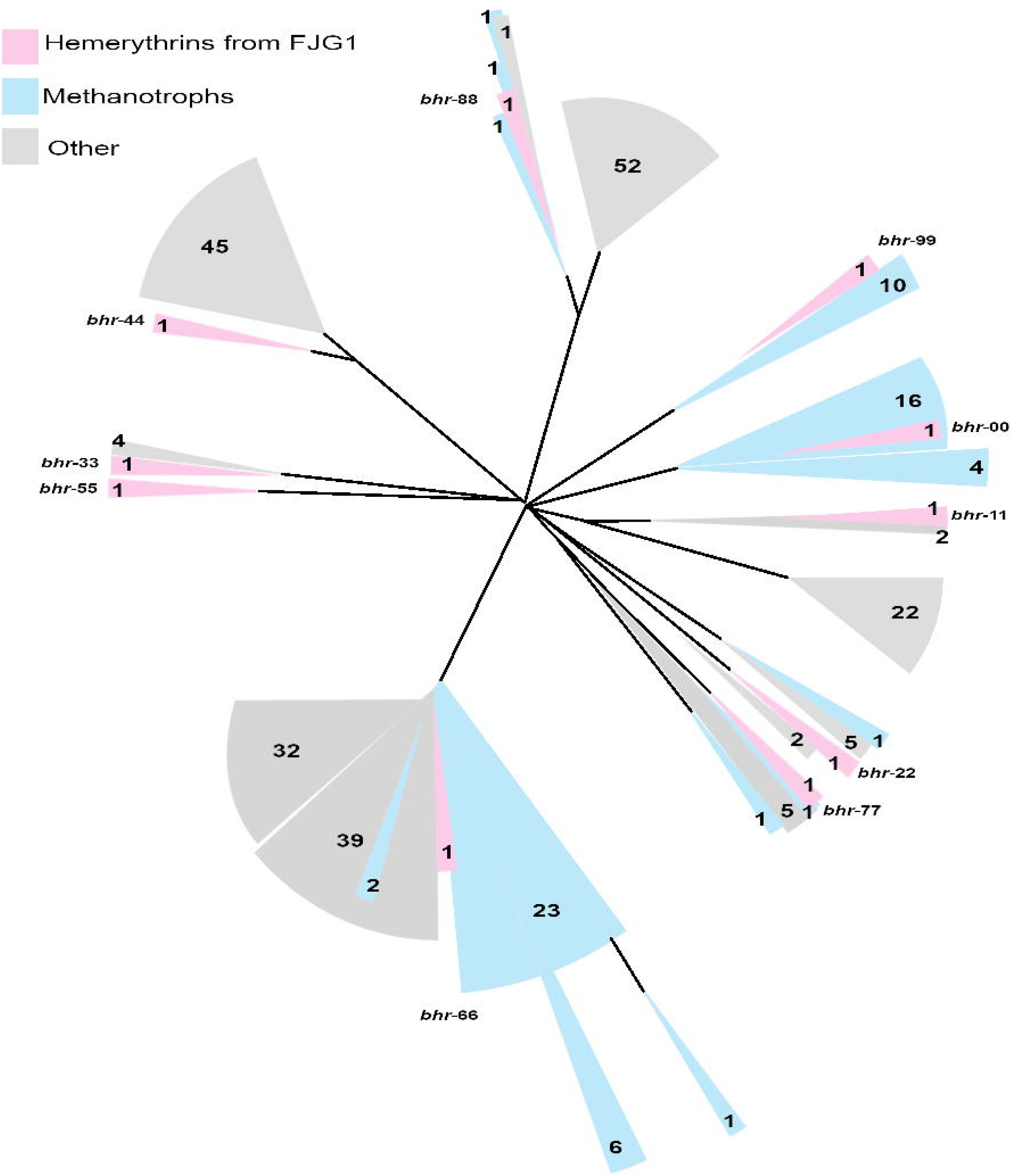
Unroooted phylogram of prokaryotic genes related to the 10 *bhr* genes from *M. denitrificans* FJG1. Homologs were identified from the GenBank database using BLASTn (Geneious Prime 2019 v1.2.3). Sequences from *M. denitrificans* FJG1 are highlighted in pink and genes from other proteobacterial methanotrophs are highlighted in blue. Non-methanotrophs are colored grey.

**Table 1.** Hemerythrin genes identified in the *Methylomonas denitrificans* FJG1 genome (CP014476)

| Name | Locus_tag/<br>GI_number | Min | Max | Length | Direction | Annotation | Protein_id | Protein | GO_function |
| --- | --- | --- | --- | --- | --- | --- | --- | --- | --- |
| *bhr-00* | JT25_RS00380<br>GI:998900537 | 85767 | 85372 | 396 | R | hemerythrin family protein | WP_036277397.1 | bacteriohemerythrin | GO:0046872 - metal ion binding |
| *bhr-11* | JT25_RS03015 | 663113 | 663445 | 333 | F | hemerythrin domain-containing protein | WP_132325259.1 | hemerythrin domain-containing protein | - |
| *bhr-22* | JT25_RS04600 | 989403 | 989813 | 411 | F | bacteriohemerythrin | WP_036273626.1 | bacteriohemerythrin | GO:0046872 - metal ion binding |
| *bhr-33* | JT25_RS05590 | 1204483 | 1204097 | 387 | R | hemerythrin family protein | WP_036274438.1 | hemerythrin family protein | - |
| *bhr-44* | JT25_RS10375 | 2251509 | 2251123 | 387 | R | hemerythrin family protein | WP_036280274.1 | hemerythrin family protein | - |
| *bhr-55* | JT25_RS11125<br>GI:1011416966 | 2420834 | 2416305 | 4530 | R | bacteriohemerythrin | WP_062328608.1 | bacteriohemerythrin | GO:0046872 - metal ion binding |
| *bhr-66* | JT25_RS17815 | 3859160 | 3860944 | 1785 | F | bacteriohemerythrin | WP_062329310.1 | bacteriohemerythrin | GO:0046872 - metal ion binding |
| *bhr-77* | JT25_RS18710 | 4064071 | 4063667 | 405 | R | bacteriohemerythrin | WP_036274237.1 | bacteriohemerythrin | GO:0046872 - metal ion binding |
| *bhr-88* | JT25_RS18715 | 4064508 | 4064095 | 414 | R | hemerythrin family protein | WP_036274239.1 | bacteriohemerythrin | GO:0046872 - metal ion binding |
| *bhr-99* | JT25_RS21450 | 4674029 | 4673625 | 405 | R | hemerythrin family protein | WP_052142344.1 | hemerythrin family protein | - |

As *bhr-*66 was the most common homolog found in bacterial genomes, we examined its detailed phylogeny with organisms for which full genomes were available (Fig. 2; Supp. Data, Sheet 4). A cladogram using full-length *bhr*-66 alignments showed its polyphyletic distribution, several instances of horizontal gene transfer, and a range of annotations (e.g. histidine kinase, response regulator, hypothetical protein) that suggest a relationship to the Bhr class of oxygen-sensing and signal transducing types (18). Expectedly, *bhr*-66 was monophyletic within the methanotrophic bacteria. We compared the phylogeny of the methanotroph *bhr*-66 genes with their 16S rRNA genes and found that they were highly congruent, suggesting an ancestral vertical transmission (Fig. 3). Similar congruence was found for the methanotroph-specific *bhr*-00 gene. When comparing phylogenies between *bhr*-00 and *bhr*-99, their high congruence and sequence similarity to each other suggests that they are paralogs (Fig. 3). Similarly, two pairs: *bhr*-33/-44, and *bhr*-77/-88, share high sequence similarity to one another, suggesting that each of the two pairs may also be paralogs (Supplemental data sheet 3). The *bhr*-77/-88 genes are encoded contiguously, perhaps due to a recent gene duplication event, and thus share the same gene neighbourhood. As very few genomes of methanotrophs or other bacteria were found to encode *bhr*-11, -22, -33, -44, -77 and -88, we could not perform a detailed phylogenetic analysis.

**Figure 2.**
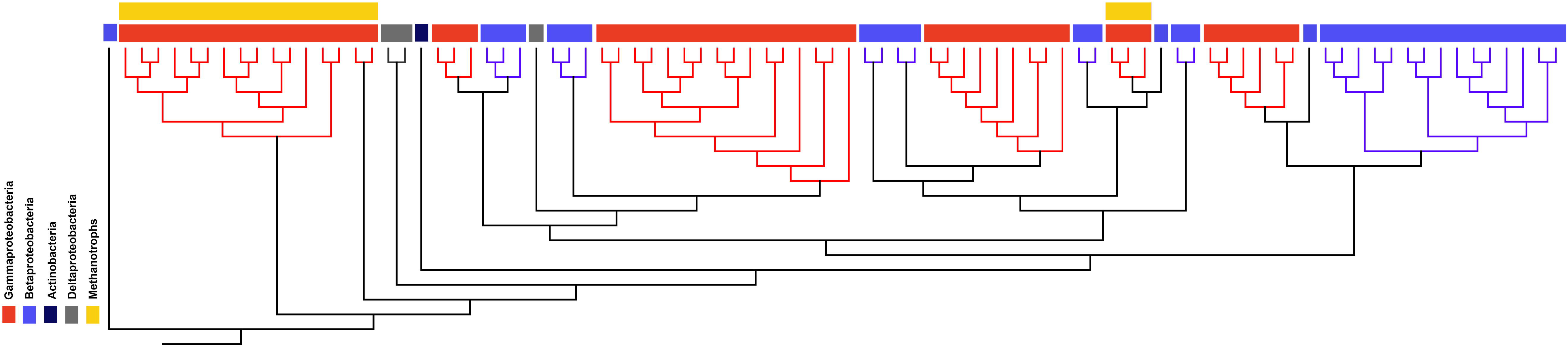
Cladogram created with full-length CDSs matching the *bhr-*66 gene of *M. denitrificans* FJG1 (rooted with eukaryotic outgroup *Themiste pyroides* (KY007473 – KY007479; Geneious Prime 2019 v1.2.3). Green: *Deltaproteobacteria*, blue: *Betaproteobacteria*, red: *Gammaproteobacteria*, purple: *Actinobacteria*. Genes from methanotrophic organisms are highlighted with a yellow bar.

**Figure 3.**
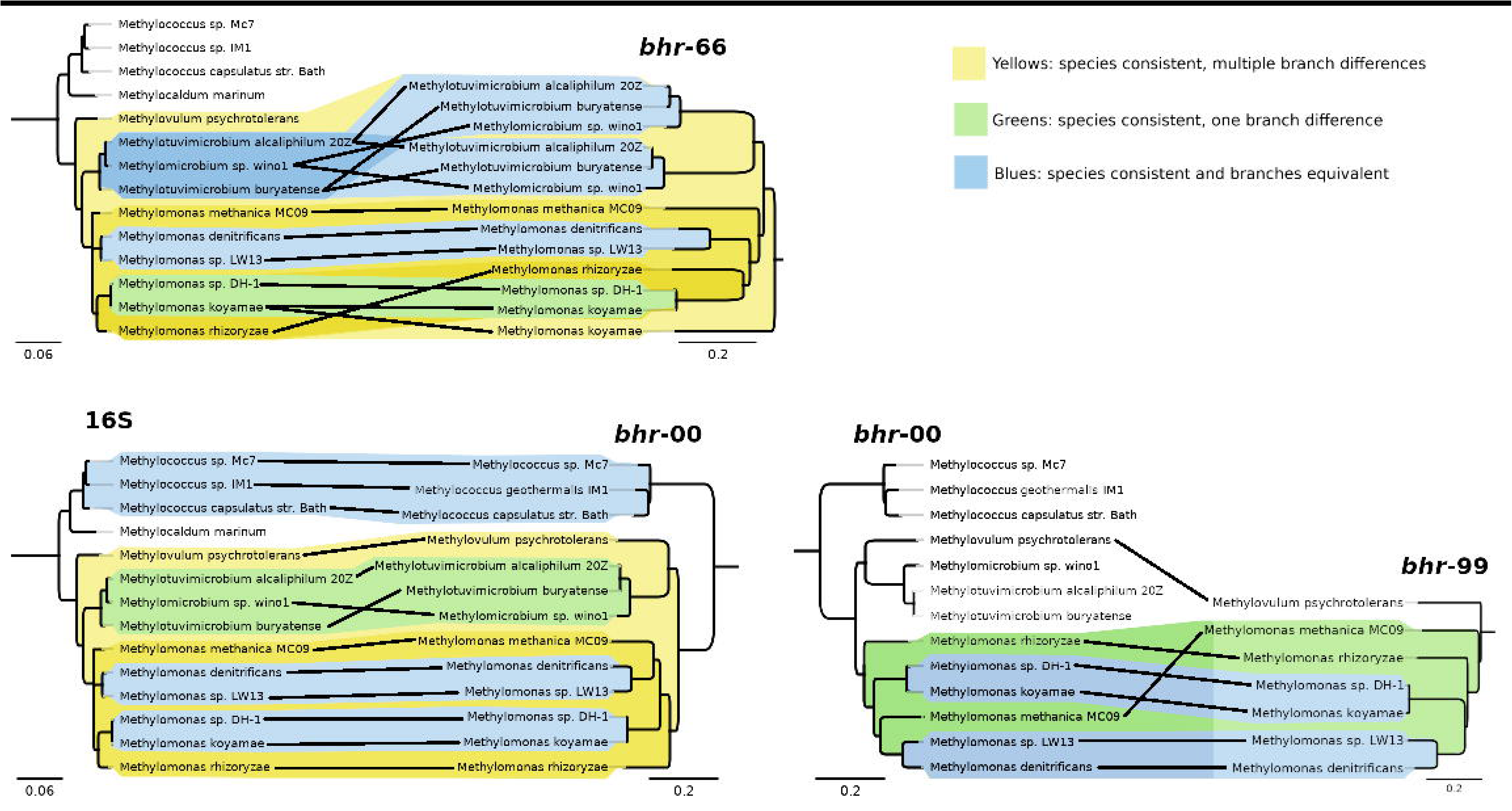
Cladograms of phylogenetic pairings between *bhr*-00 and *bhr*-66 with 16S rRNA genes in methanotrophs and of *bhr-*00 with *bhr-*99 genes (Geneious Prime 2019 v1.2.3). Yellow highlights indicate multiple branch differences, green highlights indicate one branch difference, and blue highlights indicate consistent branch topology.

### Conservation and predicted functions of *bhr* gene neighbourhoods in methanotrophic bacteria

To explore the possible functionalities of commonly shared *bhr* genes among methanotrophic bacteria, we examined the surrounding gene neighbourhoods for *bhr*-00, -66 and -99 across available complete methanotroph genomes (Fig. 4). The *bhr*-00 gene neighbourhood is highly conserved in the *Methylomonas* and *Methylotuvimicrobium* genera with flanking gene clusters for biotin synthesis (*bio*BFHCD), an enzyme co-factor, and cytochrome *c* oxidase genes (*cox*/*cta*) for oxygen respiration. The *bhr*-66 gene neighbourhood is less well conserved, with genes related to signal transduction (i.e. response regulator) and ATP-binding domains flanking *bhr*-66 in most *Methylomonas* genomes, and genes for the electron carrying cytochrome P-460 flanking *bhr*-66 in *Methylmicrobium* and *Methylotuvimicrobium* genomes (Fig. 4). Although *bhr*-99 appears to be a paralog of *bhr*-00, its gene neighbourhood is distinct and conserved in four of the *Methylomonas* genomes (Fig. 4) which show the AlkB DNA-repair gene encoded on the same DNA strand as *bhr-*99. An operon for the Na^+^-translocating NADH:quinone oxidoreductase (NQR) complex, which is involved in creating sodium motive force for transporter and flagellar function (19), is encoded on the opposing strand. The upstream region of *bhr*-99 is not well conserved among the analyzed methanotroph genomes (Fig. 4).

**Figure 4.**
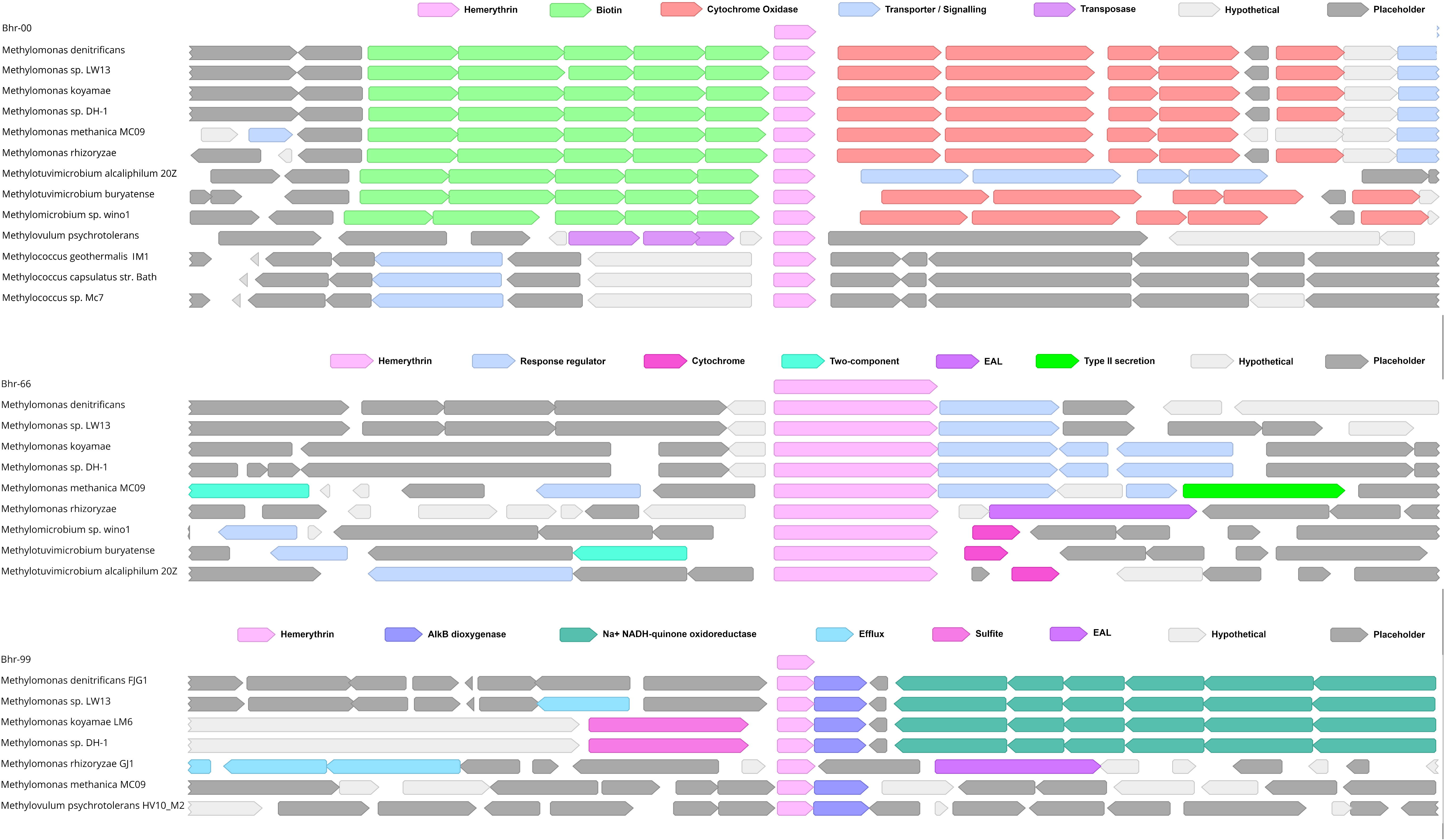
Gene neighborhoods and colour-coded gene annotations surrounding *bhr*-00, *bhr*-66, and *bhr*-99 (Geneious Prime 2019 v1.2.3). Key genes were colour-coded to discriminate functional categories as shown above each gene neighborhood.

### Expression of *bhr* homologs and their neighboring genes in *M. denitrificans* FJG1 transitioning from high to low oxygen

Expression of the *bhr* homologs and their surrounding gene neighbourhoods under oxic (24 h) and hypoxic (48 and 72 h) conditions were determined for cells grown in ammonium (AMS) or nitrate (NMS) mineral salts media (14). Among the *bhr* genes, only *bhr*-00 showed a strong (>2 norm log2 ratio) increase in mRNA abundance under hypoxia in both NMS and AMS media (Fig. 5, Table 2, Supp. Fig. 3). This transcript was also, by orders of magnitude, the most highly expressed among the *bhr* homologs (Fig. 5, Table 2). Even though previous research showed that *M. denitrificans* FJG1 reduces nitrate only when grown in NMS (14), the AMS-grown bacteria also expressed *bhr*-00 mRNA to the same magnitude under hypoxia (Fig. 5, Table 2), indicating that oxygen deprivation and not N-source was the signal affecting its expression. Other *bhr* homologs (*bhr*-11, -55, -66, -77, -88) showed significantly increased mRNA expression (P<0.05) under hypoxia (ratio of 24:48 h time points) when *M. denitrificans* FJG1 was grown in NMS, but not in AMS, media suggesting a potential linkage to denitrification functions (Table 2). Proteomic analysis was performed only for NMS-grown cells to compare expression responses at the protein and mRNA levels. We detected both Bhr-00 and Bhr-55 in the proteome (accession numbers GI:998900537, GI:1011416966 respectively) (Fig. 5, Supp. Fig. 3, Supp. Data sheet 6). Bhr-00 increased by around 6-fold from the oxic to the hypoxic condition, making up >1% of the total proteome. In contrast, Bhr-55 was of much lower abundance (<0.001%) and it decreased from the oxic to hypoxic condition, contrary to expression of its transcript.

**Figure 5.**
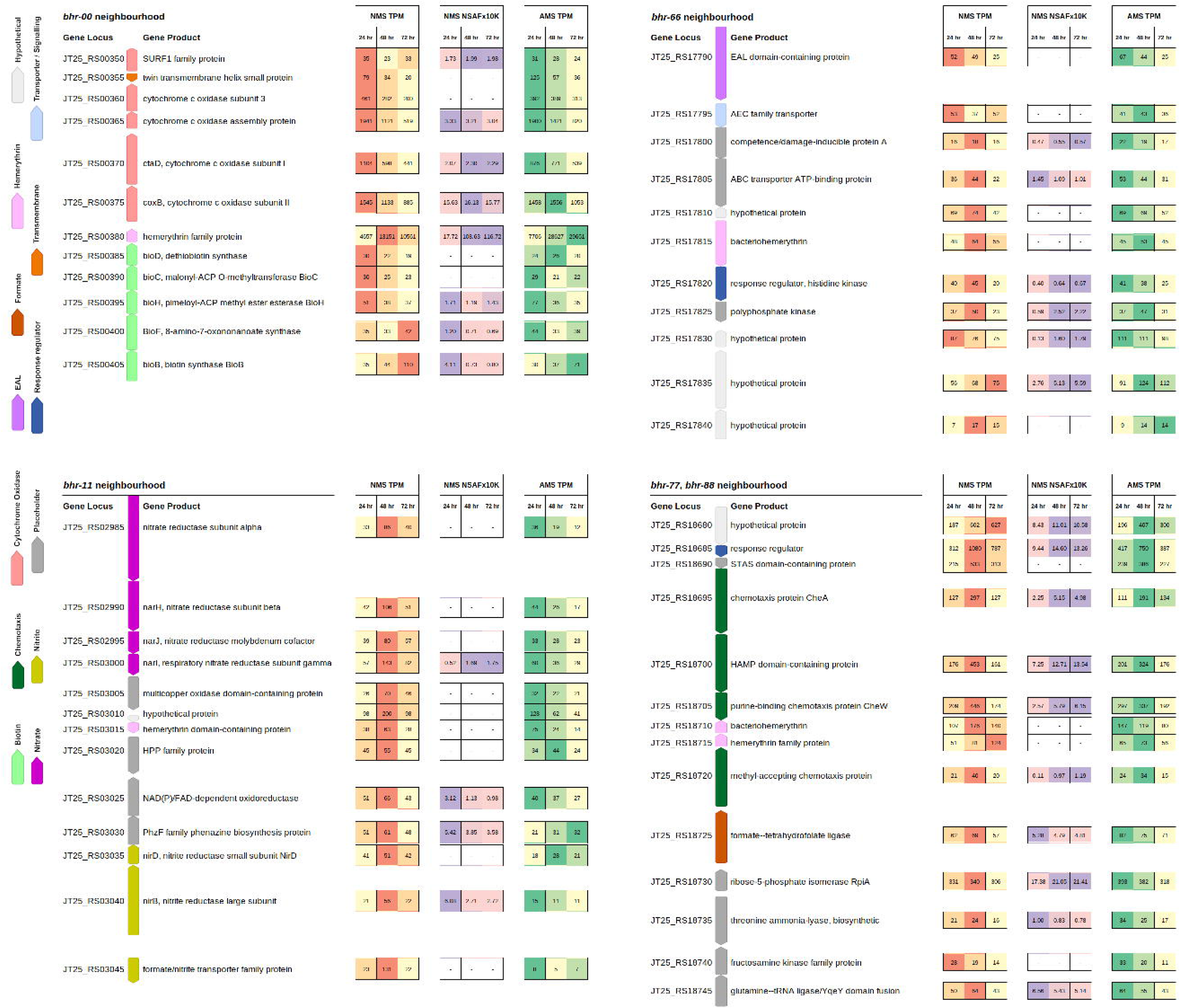
Gene neighborhoods of *bhr-*00, -11, -66, and -77/-88 with gene expression and protein abundance data. Gene neighborhoods are shown on the left in their operon structures and are colour-coded by functional group as per the legend. Up arrows indicate genes encoded on the forward DNA strand, and down arrows indicate genes encoded on the reverse DNA strand. Columnar data displayed to the right represents mRNA and protein expression abundance for *M. denitrificans* FJG1 cells transitioning from oxic (24 h) through hypoxic (48 h) to anoxic (72 h) conditions in rows from left to right as noted in the column headings. mRNA expression (TPM) in NMS: low (yellow) to high (red) and AMS: low (yellow) to high (green). Protein expression (NSAF) for NMS: low (pink) to high (purple). Rows are aligned with the center point of corresponding gene icons in the neighborhoods to the left. CDS lengths are proportional and gaps are preserved. (Neighborhood genes colourized in Geneious Prime 2019 v1.2.3. Data columns prepared in LibreOffice Calc. Components organized and legend added in GIMP 2.10.)

**Table 2.** Differential expression of *bhr* gene homologues from *M. denitrificans* FJG1 based on RNAseq data collected after 24 and 48 h growth in NMS or AMS media with methane.

|  | Nitrate Treatment (NMS) |  |  |  |  |
| --- | --- | --- | --- | --- | --- |
|  | Raw Expression (TPM) |  | Differential Expression<br>24h vs 48h |  |  |
|  | 24h | 48h | log2 ratio | adjusted<br>p-value | normalized<br>log2 ratio |
| *bhr-00* | 4,657.50 | 13,150.82 | 1.35 | 0.000 | 3.33 |
| *bhr-11* | 38.22 | 62.58 | 0.47 | 0.034 | 2.45 |
| *bhr-22* | 93.16 | 61.85 | -0.65 | 0.000 | 1.33 |
| *bhr-33* | 29.11 | 41.03 | 0.50 | 0.051 | 2.48 |
| *bhr-44* | 83.43 | 72.20 | -0.28 | 0.104 | 1.70 |
| *bhr-55* | 30.82 | 46.69 | 0.41 | 0.000 | 2.39 |
| *bhr-66* | 48.44 | 63.83 | 0.28 | 0.002 | 2.26 |
| *bhr-77* | 107.16 | 176.26 | 0.62 | 0.000 | 2.60 |
| *bhr-88* | 50.66 | 80.73 | 0.59 | 0.000 | 2.57 |
| *bhr-99* | 170.36 | 172.87 | -0.05 | 0.732 | 1.93 |
| *rpoB* | 418.02 | 113.34 | -1.98 | 0.000 | - |

|  | Ammonium Treatment (AMS) |  |  |  |  |
| --- | --- | --- | --- | --- | --- |
|  | Raw Expression (TPM) |  | Differential Expression<br>24h vs 48h |  |  |
|  | 24h | 48h | log2 ratio | adjusted<br>p-value | normalized<br>log2 ratio |
| *bhr-00* | 7,706.16 | 28,626.86 | 1.91 | 0.000 | 3.21 |
| *bhr-11* | 75.47 | 23.55 | -1.14 | 0.000 | 0.16 |
| *bhr-22* | 122.98 | 57.33 | -0.57 | 0.000 | 0.73 |
| *bhr-33* | 22.58 | 39.44 | 0.50 | 0.004 | 1.80 |
| *bhr-44* | 74.32 | 65.09 | -0.45 | 0.000 | 0.85 |
| *bhr-55* | 32.34 | 36.47 | 0.25 | 0.000 | 1.55 |
| *bhr-66* | 45.44 | 53.34 | 0.17 | 0.013 | 1.47 |
| *bhr-77* | 146.80 | 119.38 | 0.07 | 0.470 | 1.37 |
| *bhr-88* | 64.90 | 73.10 | 0.22 | 0.071 | 1.52 |
| *bhr-99* | 193.38 | 181.93 | -0.15 | 0.036 | 1.15 |
| *rpoB* | 421.34 | 162.99 | -1.30 | 0.000 | - |

While genes for cytochrome *c* oxidase in the gene neighborhood of *bhr*-00 showed decreased expression at the mRNA level under hypoxia, the protein abundances remained relatively constant for this gene cluster (Fig. 5). In the same gene neighborhood, the *bioB* biotin synthase gene showed decreased mRNA, but increased protein abundance with hypoxia. Genes encoding the enzymes for nitrate reduction within the gene neighbourhood of *bhr*-11 showed increased mRNA levels under hypoxia for only NMS-grown cells, which is consistent with the onset of denitrification activity (14) (Fig. 5). Only the NarI protein in the nitrate reductase gene cluster was detected in the proteome, but its expression was elevated under hypoxia. The gene neighbourhood of *bhr*-66 showed increased expression at the mRNA and protein levels for the response regulator histidine kinase and polyphosphate kinase as NMS-grown cells transitioned from oxic to hypoxic conditions, suggesting potential involvement in oxygen sensing under denitrifying conditions (Fig. 5). Furthermore, the gene neighbourhood surrounding *bhr*-77/-88 includes multiple genes for chemotaxis that showed increased expression at both the mRNA and protein levels in NMS-grown cells as they transitioned from oxic to hypoxic conditions, in the same range as the nitrate reduction genes (Fig. 5, Supp. Table 1). The results suggest that chemotaxis and oxygen sensing may be facilitated by *bhr*-domain proteins in *M. denitrificans*, which is consistent with Bhr functions characterized in other bacteria (7, 18, 20).

### Diversity of Bhr protein structures from *M. denitrificans* FJG1

Because the 10 *bhr* genes of *M. denitrificans* FJG1 vary in size and genomic context, we explored their predicted protein structural diversity using AlphaFold2 (Fig. 6A). Each predicted structure showed unique features, although the Bhr-33 and -44 structures were similar to one another. When the predicted structure of Bhr-00 was overlaid with the experimentally derived crystal structure of the Bhr-Bath protein, which was demonstrated as an oxygen binding and delivery system to pMMO enzymes in cell-free membrane systems (11), they were congruent even though their amino acid identities overlapped by only 57.58% (Fig. 6B, Supp. Fig. S2, Supp. Movie).

**Figure 6.**
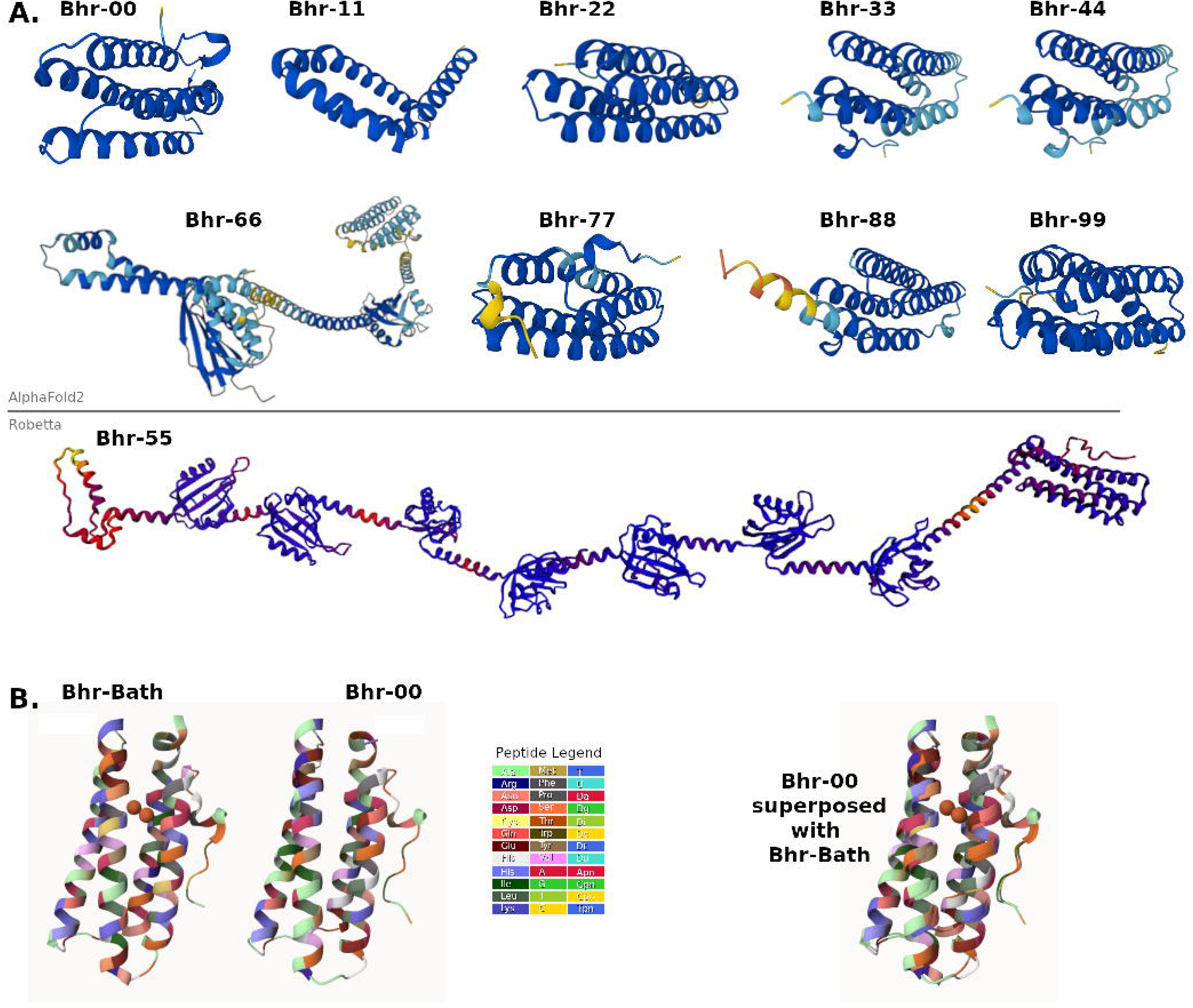
AlphaFold2 Predicted Protein Structures visualized in Mol* Viewer. **A.** Bhr protein structures predicted from *M. denitrificans* FJG1 genes from the AlphaFold2 database. Br-55 was not found using AlphaFold2 and was instead created using Robetta, as described in Methods. **B.** Structural comparison of predicted AlphaFold2 structure for Bhr-00 (ID: A0A140E3I1) with experimentally determined Bhr-Bath structure (Chen et al. 2015, PDB ID: 4XPX). Diiron cofactor is shown in orange. Animated superposition in .mp4 format is available as a Supplemental Movie.

The hybrid structure of the Bhr-66 protein of *M. denitrificans* FJG1explains the inconsistency in its gene annotation and phylogeny. This predicted structure has a distinct bacteriohemerythrin of four alpha-helices arranged in a chimney shape at the N-terminus, which was the segment likely identified as a bacteriohemerythrin by nBLAST, appended by a linear C-terminal extension containing two structures with distinctive folds reminiscent of PAS domains of s-box histidine kinases and response regulators. The overall structure suggests that Bhr-66 acts as part of an oxygen-sensing two-component regulatory system that has been well described previously in other bacteria (1, 21, 22). This may also explain the proximity of genes grouped as response regulators in the neighbourhoods of several methanotroph *bhr*-66 genes (Fig. 4).

Similarly, the predicted Bhr-55 structure appears to have a bacteriohemerythrin segment at its N-terminus with a long trailing section at the C-terminus, which includes up to seven folded PAS-like domains (Fig. 6). Interestingly, no sequences related to *bhr*-55 were identified in other genomes, suggesting that this protein may be unique to *M. denitrificans* FJG1.

## Discussion

*Methylomonas denitrificans* FJG1 consumes methane and reduces nitrate to survive hypoxia (14). Close examination of the *M. denitrificans* FJG1 genome revealed 10 homologs of *bhr* genes, suggesting multiple potential functions associated with oxygen sensing and binding in this bacterium. By providing a detailed examination of the phylogeny, gene neighbourhood, expression, and predicted protein structures of the 10 *bhr* genes of *M. denitrificans* FJG1, we provide compelling evidence that Bhr-00, a bacteriohemerythrin that appears nearly exclusive to gammaproteobacterial methanotrophs, is likely the oxygen-binding protein responsible for supporting methane oxidation in low oxygen environments. This assumption is partially based on a close match of the predicted structure of Bhr-00 to Bhr-Bath, which was shown to bind and deliver oxygen to pMMO enzymes *in vitro* (11). Elevated expression of *bhr*-00 at both the transcript and protein levels in *M. denitrificans* FJG1 under hypoxic conditions further supports this hypothesis. Also in support, a study of *Methylomonas* sp. LW13 with a deleted *bhr*-00 gene was unable to grow in methane-oxygen counter gradient, but was able to grow normally in planktonic culture with 50% methane:50% air (13). Aside from Bhr-00, this study identified *bhr* homologs co-localized with denitrification (*bhr*-11), oxygen sensing (*bhr*-66), and chemotaxis (*bhr*-77, -88) associated genes. These co-localized gene clusters showed increased mRNA expression under hypoxic conditions with NMS, suggesting multiple roles for *bhr* domain-containing proteins in *M. denitrificans* FJG1 for its survival and respiration under low oxygen, denitrifying, conditions.

The hemerythrin originally discovered in marine worms, Hr, is encoded by a ∼400 bp gene for a monomeric protein that assembles into an octameric oxygen transport protein, causing the pink hue of the worms’ blood (2). The Bhr-Bath protein purified from *M. capsulatus* Bath was crystallized as a monomer of similar size and shape to Hr and was shown to enhance oxygen delivery from the cytoplasm to pMMO-enriched membrane fractions (9, 10). Purified Hr and Bhr-Bath proteins turn an opaque reddish-purple or bright pink colour when bound to oxygen (10, 23). This is the same hue as *M. denitrificans* FJG1 cells when grown to stationary phase in liquid culture or on plates, further suggesting that increased expression of Bhr could be responsible for oxygen binding and thus the change in their coloration beyond that of orange-hued carotenoid proteins (24).

Aside from the identification and proposed role of Bhr-00, this study revealed several diverse *bhr* gene homologs encoded by gammaproteobacterial methanotrophs. Based on our findings, relevant potential functions of *bhr*-containing proteins in methanotrophs could include oxygen binding and delivery to support methane oxidation and respiration, oxygen sensing, nitrate reduction, and chemotaxis. Gammaproteobacterial methanotrophs often in the *Methylomonas* genus are found in hypoxic to anoxic ecosystems, particularly in freshwater lakes (25). The discovery and hypoxic regulation of the methanotroph-specific *bhr*-00 gene suggests a mechanism to promote the binding and delivery of oxygen to support methane oxidation. Moreover, the apparent restriction of *bhr*-00 to only a few related genera (*Methylomonas*, *Methylotuvimicrobium*, *Methylomicrobium*) could explain their enrichment in these ecosystems (25). From an environmental perspective, methanotrophs play a key role in biogeochemical cycling of methane and nitrogen (26). Methane saturation of nutrient-impacted lakes and reservoirs may further enrich for methanotrophs that rely on *bhr* to survive and thrive as these ecosystems transition into hypoxia. The emergence of methanotroph phylotypes similar to *M. denitrificans* FJG1 should be carefully monitored in nutrient impacted freshwater ecosystems as they are bellwethers for hypoxia, methane cycling, and N_2_O production (25, 27).

## Materials and Methods

### Phylogenetic analyses

The genome sequence of *M. denitrificans* FJGI (GenBank accession CP014476A) was downloaded and analyzed using Geneious Prime 2019 (v1.2.3). The FJG1 genome was found to contain 10 coding sequences (CDS) annotated as hemerythrin, bacteriohemerythrin, hemerythrin family, or hemerythrin domain-containing proteins in the GenBank and RefSeq annotation databases (Table 1). To facilitate further analyses, the 10 sequences were designated *bhr*-00 through *bhr*-99 in the order of location from the genome origin of replication for the purposes of this study.

Unrooted phylogrammatic overview of all *M. denitrificans* FJG1 *bhr* BLASTn hits in prokaryotes.

Phylogenetic analyses of the 10 *bhr* sequences from *M. denitrificans* FJG1 were performed using Geneious Prime 2019 (v1.2.3). The BLASTn plug-in was used with default settings. The resulting dataset was filtered by excluding eukaryotic organisms and prokaryotic organisms containing fewer than two *bhr* BLASTn hits.) Because the bhr-66 sequence is so widespread among prokaryotes of multiple lineages, we chose to focus on a smaller dataset including organisms that contain not just the ubiquitous *bhr*-66 but multiple hemerythrin sequences that are more differentiated.

The resulting BLASTn hits were collected into a database for analysis and categorized by aligning them to a concatenated sequence consisting of the 10 FJG1 genes. The reference sequence was created by extracting the 10 *bhr* genes from the FJG1 genome and concatenating them in order, separated by spacers of 100 base pairs. The concatenated reference sequence eliminates duplicate matches and ensures that BLASTn hits are correlated to the FJG1 numbered sequence with which they share the greatest sequence similarity (Supp. Data, Sheet 3). After aligning all of the selected prokaryotic BLASTn hits with the reference sequence, an unrooted phylogram was generated showing the inclusive overview of a broad range of *bhr* genes among prokaryotic organisms (Geneious Prime 1.2.3, Global alignment with free end gaps, Cost matrix 51% similarity (5.0/-3.0), Tamura-Nei, Neighbor-Joining (NJ), No Outgroup, Gap open penalty 12, Gap extension penalty 3, Automatically determine direction). The accompanying data file of sequence similarity presented in a distance matrix is available as Supp. Data, Sheet 3.

The BLASTn hits were used to locate prokaryotic organisms containing putative *bhr* sequences with available complete genomes. These *bhr*-containing prokaryote genomes were downloaded from GenBank and assembled into a local genome database. A circular alignment of methanotroph genomes containing *bhr* sequences was generated using BLAST Ring Image Generator (BRIGv0.95 default settings, Upper threshold 70%, Lower threshold 50%) to show the location of their *bhr* sequences in relation to each other (Accession numbers CP014476, CP033381, CP023669, CP002738, CP014360, CP022129, CP035467, CP024202, CP046565, FO082060, NC_002977.6, AP017928). The circular alignment is available as Supp. Fig. S1.

Working in the locally assembled *bhr*-containing genome database, the *bhr* BLAST hit sequences were used to locate full length CDSs in all collected genomes. These full sequences including start and stop codons were extracted and compiled into a comprehensive *bhr* CDS database to facilitate more accurate and complete phylogenetic analyses. Sequences were first aligned using the Geneious Alignment tool (Global Alignment with free end gaps, Cost Matrix: 51%, default settings). Based on these alignments, three rooted cladograms were generated using Geneious Prime 2019, one for each of the *M. denitrificans* FJG1 genes. CDSs with four or more matching organisms (total n ≥ 5) (Geneious Tree Builder v1.2.3, Genetic Distance Model: Tamura-Nei, Tree Build Method: Neighbour-joining, Resampling Method: 1,000 bootstrap, Support Threshold: 50%, Image Scale: Substitutions per site). Less than four genomes could be located encoding *bhr-*11, -22, -33, -44, -77 and/or -88, and therefore we could not perform a detailed analysis of these genes across bacterial or other methanotroph species.

Cladograms describing *bhr* sequences *bhr*-00 and *bhr*-99 from methanotrophs were rooted with the prokaryotic outlier *Chitinomonas* sp R3-44 (CP041730); however, in order to include all prokaryotes for the widely distributed *bhr*-66 sequence, the cladogram was rooted using hemerythrin sequences from a eukaryote outgroup, *Themiste pyroides* (KY007473.1, KY007474.1, KY007475.1, KY007476.1, KY007477.1, KY007478.1, KY007479.1). For phylogenetic comparison, we also generated 16S rRNA trees using the same methods. Comparative images were created using Geneious Prime 2019 and colour was added to highlight branch differences.

### Identification and comparison of *bhr* gene neighbourhoods

Three of the 10 *bhr* sequences (*bhr*-00, *bhr*-66, and *bhr*-99) were selected for gene neighbourhood analysis, as at least five methanotroph genomes contained homologous copies to those found in *M. denitrificans* FJG1. Less than four genomes encoding *bhr*-11, -22, -33, -44, -77 and/or -88, could be located therefore we could not perform a meaningful comparative analysis of these gene neighbourhoods. Approximately 16,000 bp surrounding each *bhr* sequence was extracted from each annotated genome. In cases where multiple genomes were found for a species, the most recent annotated genome assembly was used to leverage the most up-to-date annotations available. In cases where full genomes were found, but lacked gene annotations, they were annotated in-house using Geneious Prime 2019 (Default settings, Similarity > 50%), drawing from the in-house database of all annotated methylotroph genomes downloaded for this project (> 2,000 genomes). Comparative genomics based on these neighbourhood extractions was used to investigate each *bhr* sequence. Annotations were compared manually to overcome inconsistencies in labeling methodology. Colour schemes were used to identify sequences sharing similar or related functional categories, while maintaining the annotation labeling of the original authors. An accompanying comparison of the *bhr*-66 gene neighbourhood in non-methanotroph genomes is available as Supp. Fig. S4.

### RNAseq analysis

The RNAseq experiments were performed as described previously (14). Briefly, *M. denitrificans* FJG1 was cultured in NMS or AMS medium at 30% methane to 70% air to allow for natural oxygen depletion over time. The cultures achieved hypoxia and entry into stationary phase by 48 h (Supp. Fig. S5). Replicate cultures (n=5) were destructively sampled at 24 and 48 h time points to determine differential mRNA expression as a function of oxygen depletion. Cells were collected and mRNA was purified, converted to cDNA, and processed for transcriptome analysis using Ilumina Hi-Seq 2000 sequencing technology as described (13). Geneious Prime 2019 (v1.2.3) was employed with the DESeq2 plug-in (default modelsettings “Parametric”) to re-calculate expression values and differential expression (14). PCA plots revealed zero outliers with < 7% total within-treatment variance. TPM values for transcripts at 24, 48 and 72 h time points for *bhr* homologs and members of their gene neighborhoods were calculated using DESeq2 in Geneious Prime 2019 (v1.2.3), and the ratio of differential expression of *bhr* homologs between 24 and 48 h were determined using R.

### Protein extraction and peptide preparation

For each of the three treatments we prepared tryptic digests from six biological replicates following the filter-aided sample preparation (FASP) protocol described by Wisniewski et al. (28). SDT-lysis buffer (4% (w/v) SDS, 100 mM Tris-HCl pH 7.6, 0.1 M DTT) was added in a 1:10 sample/buffer ratio to the sample pellets. Samples were heated for lysis to 95° C for 10 minutes followed by pelleting of debris for 5 min at 21,000 x g. 30 µl of the cleared lysate were mixed with 200 µl of UA solution (8 M urea in 0.1 M Tris/HCl pH 8.5) in a 10 kDa MWCO 500 µl centrifugal filter unit (VWR International) and centrifuged at 14,000 x g for 40 min. 200 µl of UA solution were added again and centrifugal filter spun at 14,000 x g for 40 min. 100 µl of IAA solution (0.05 M iodoacetamide in UA solution) were added to the filter and incubated at 22° C for 20 min. The IAA solution was removed by centrifugation and the filter was washed three times by adding 100 µl of UA solution and then centrifuging. The buffer on the filter was then changed to ABC (50 mM Ammonium Bicarbonate), by washing the filter three times with 100 µl of ABC. 1 µg of MS grade trypsin (Thermo Scientific Pierce, Rockford, IL, USA) in 40 µl of ABC were added to the filter and filters incubated overnight in a wet chamber at 37° C. The next day, peptides were eluted by centrifugation at 14,000 x g for 20 min, followed by addition of 50 µl of 0.5 M NaCl and again centrifugation. Peptides were desalted using C18 spin columns (Thermo Scientific Pierce, Rockford, IL, USA) according to the manufacturer’s instructions. Approximate peptide concentrations were determined using the Pierce Micro BCA assay (Thermo Scientific Pierce, Rockford, IL, USA) following the manufacturer’s instructions.

### Proteomics 1D-LC-MS/MS

Samples were analyzed by 1D-LC-MS/MS using a block-randomized design as outlined by Oberg and Vitek (29). Two blank runs were done between samples to reduce carry over. For each run 800 ng of peptide were loaded onto a 2 cm, 75 µm ID C18 Acclaim® PepMap 100 pre-column (Thermo Fisher Scientific) using an EASY-nLC 1000 Liquid Chromatograph (Thermo Fisher Scientific) set up in 2-column mode. The pre-column was connected to a 50 cm x 75 µm analytical EASY-Spray column packed with PepMap RSLC C18, 2µm material (Thermo Fisher Scientific), which was heated to 35° C via the integrated heating module. The analytical column was connected via an Easy-Spray source to a Q Exactive Plus hybrid quadrupole-Orbitrap mass spectrometer (Thermo Fisher Scientific). Peptides were separated on the analytical column at a flow rate of 225 nl/min using a 260 min gradient going from buffer A (0.2% formic acid, 5% acetonitrile) to 20% buffer B (0.2% formic acid in acetonitrile) in 200 min, then from 20 to 35% B in 40 min and ending with 20 min at 100% B. Eluting peptides were ionized with electrospray ionization (ESI) and analyzed in the Q Exactive Plus. Full scans were acquired in the Orbitrap at 70,000 resolution. MS/MS scans of the 15 most abundant precursor ions were acquired in the Orbitrap at 17,500 resolution. The mass (m/z) 445.12003 was used as lock mass. Ions with charge state +1 were excluded from MS/MS analysis. Dynamic exclusion was set to 30 sec. Roughly 160,000 MS/MS spectra were acquired per sample run.

### Protein identification, quantification and statistics

For protein identification a database was created using all protein-coding gene sequences from the complete genome of *Methylomonas denitrificans* strain FJG1 (Genbank Identifier CP014476.1). The database was submitted to the PRIDE repository (see below). For protein identification MS/MS spectra were searched against the database using the Sequest HT node in Proteome Discoverer version 2.0.0.802 (Thermo Fisher Scientific) with the following parameters: Trypsin (Full), max. 2 missed cleavages, 10 ppm precursor mass tolerance, 0.1 Da fragment mass tolerance and max. 3 equal dynamic modifications per peptide. The following three dynamic modifications were considered: oxidation on M (+15.995 Da), carbamidomethyl on C (+57.021 Da) and acetyl on protein N-terminus (+42.011 Da). False discovery rates (FDRs) for peptide spectral matches (PSMs) were calculated and filtered using the Percolator Node in Proteome Discoverer (30). Percolator was run with the following settings: Maximum Delta Cn 0.05, a strict target FDR of 0.01, a relaxed target FDR of 0.05 and validation based on q-value. Search results for all 18 samples were combined into a multiconsensus report using the FidoCT node in Proteome Discoverer to restrict the protein-level FDR to below 5% (FidoCT q-value <0.05)(31). Based on these filtering criteria a total of 1833 proteins were identified in all samples together.

For protein quantification normalized spectral abundance factors (NSAFs) were calculated based on number of PSMs per protein using the method described by Florens et al. (32) and multiplied by 10,000. The NSAFx10,000 gives the relative abundance of a protein in a sample as a fraction of 10,000. The table with NSAFx10,000 values was loaded into the Perseus software (version 1.5.1.6, http://www.perseus-framework.org/doku.php) (33) an annotation row was added to group replicates by treatment and proteins that did not have abundance values >0 for all replicates in at least one treatment group were removed (1831 proteins remained). NSAFx10,000 values were log2 transformed. Missing values produced by log2(0) were replaced by sampling from a normal distribution assuming that the missing values are on the lower end of abundance (normal distribution parameters in Perseus: width 0.3, down shift 1.8, do separately for each column). A t-test with permutation-based FDR calculation to account for the multiple hypothesis testing problem was used to detect proteins that differed significantly in their abundance level between two treatments. The following parameters were used for the test: groupings were not preserved for randomizations, both sides, 250 randomizations, FDR of 1% and s0 of 0.

### Protein Structure Predictions

Predicted protein structures corresponding to the *bhr* gene loci for *M. denitrificans* FJG1 were downloaded from AlphaFold2 (alphafold.ebi.ac.uk) (34, 35). Nine structures were predicted that corresponded to Bhr-00 (A0A140E3I1), Bhr-11 (A0A140E4Z5), Bhr-22 (A0A126T104), Bhr-33 (A0A126T1L7), Bhr-44 (A0A126T494), Bhr-66 (A0A126T8C7), Bhr-77 (A0A126T8U2), Bhr-88 (A0A126T8Y8), and Bhr-99 (A0A140E6J4). The predicted structure of Bhr-66 was further investigated by extracting the hemerythrin section at the N-terminus and separating it from the C-terminal section of the gene into two unique nucleotide subsequences. The *bhr*-like subsequence (∼400 bp) and the C-terminal tail subsequence (∼1,200 bp) were used for BLASTn searches in the GenBank nucleotide database (Geneious Prime 2019, v1.2.3 with BLAST plugin, default settings, filtered by Grade > 40%) and the annotations from matching sequences were examined for each individual subsection of the atypical *bhr*-66 gene, revealing a hemerythrin domain at the N-terminus and a chain of multiple PAS sensing domains at the C-terminal end. No corresponding structure for Bhr-55 was available in the AlphaFold database, thus an alternative protein structure prediction tool was used (https://robetta.bakerlab.org/). The FASTA sequence of the *bhr*-55 gene was uploaded to the online tool and prediction was performed using RoseTTAFold, with default settings on the translated protein. The resulting visualization was coloured according to Error Estimate in the Robetta interface.

The predicted Bhr-00 (A0A140E3I1) structure and the Bhr-Bath structure (PDB ID: 4XPX) were downloaded from the RCSB Protein Data Bank (36) and imported into the MolStar Viewer (37) to compare their structures. Protein structures were superposed using a single Lys residue in each amino acid sequence (Lys 68, Bhr-Bath and Lys 70, Bhr-00). The Mol* animation is available as Supp. Movie. The AA alignment of FJG1 Bhr-00 and Bhr-Bath is available as Supp. Fig. S2.

## Acknowledgments

We thank Marc Strous (Univ. Calgary) for access to the proteomics equipment. This work was funded by the Canada First Research Excellence Fund Future Energy Systems to DS and LYS, an NSERC Discovery grant to LYS, and a Mitacs Accelerate grant to DS. The purchase of the proteomics equipment was supported by a grant of the Canadian Foundation for Innovation to Marc Strous. MK was supported by a NSERC Banting Postdoctoral Fellowship. The funders had no role in study design, data collection and interpretation, or the decision to submit the work for publication.

## Author Contributions

C.W., D.S. and L.Y.S. participated in the planning of the study. C.W. performed the phylogenetic analyses, comparative genomics, and drafted the manuscript. K.D.K. performed the transcriptome and proteome experiments, analyzed data, and edited the manuscript. M.K. provided the proteomic data, statistical analysis, and methodology description. D.S. and L.Y.S. provided financial support, critical analysis, writing and editing of the manuscript. All authors have read and approved the manuscript for publication.

## Originality statement

None of the work in this manuscript has been published or submitted for publication elsewhere.

## Data Availability Statement

Transcriptome data can be found in GenBank (accession number SRX696231). The genomic datasets can be found by provided accession numbers in the article. Additional details on data used in this study are in the Supp. Data file. The mass spectrometry proteomics data and the protein sequence database have been deposited to the ProteomeXchange Consortium (38) via the PRIDE partner repository with the dataset identifier PXD004041 [Reviewer access: log in at http://www.ebi.ac.uk/pride/archive/ with Username: and Password: B6gFiqkh].

## Supplemental Figure Legends

**Supp. Fig. S1**. Circular alignment of methanotroph genomes created with Blast Ring Image Generator (BRIG). The location of *bhr* genes in *M. denitrificans* FJG1 and other methanotroph genomes are designated by colored bars.

**Supp. Fig. S2.** Alignment of Bhr-00 protein sequence from *Methylomonas denitrificans* FJG1 with Bhr-Bath protein sequence from *Methylococcus capsulatus* Bath.

**Supp. Fig. S3.** Gene neighborhoods of bhr-22, -33, -44, -55 with corresponding gene expression and protein abundance data. Gene neighborhoods are shown on the left in their operon structures and are colour-coded by functional group as per the legend. Up arrows indicate genes encoded on the forward DNA strand, and down arrows indicate genes encoded on the reverse DNA strand. Columnar data displayed to the right represents mRNA and protein expression abundance for M. denitrificans FJG1 cells transitioning from oxic (24 h) through hypoxic (48 h) to anoxic (72 h) conditions in rows from left to right as noted in the column headings. mRNA expression (TPM) in NMS: low (yellow) to high (red) and AMS: low (yellow) to high (green). Protein expression (NSAF) for NMS: low (pink) to high (purple). Rows are aligned with the center point of corresponding gene icons in the neighborhoods to the left. CDS lengths are proportional and gaps are preserved. (Neighborhood genes colourized in Geneious Prime 2019 v1.2.3. Data columns prepared in LibreOffice Calc. Components organized and legend added in GIMP 2.10.)

**Supp. Fig. S4.** Gene neighborhoods of *bhr*-66 of non-methanotroph genomes.

**Supp. Fig. S5.** Growth curve of *M. denitrificans* FJG1 on NMS or AMS media. (A) Growth over time. (B) Oxygen consumption over time. Reprinted with permission from Environmental Microbiology (14).

**Supp. Table 1.** Differential expression of neighboring genes upstream and downstream of the *bhr-*00, -11, -66, and -77/-88 homologs from *M. denitrificans* FJG1 based on RNAseq data at 24 h (oxic) versus 48 h (hypoxic) growth in NMS and AMS media.

**Supp. Data.** Spreadsheet in .xls format containing metadata: Sheet 1 - Metadata; Sheet 2-list of methanotroph genes used to create rooted phylograms; Sheet 3-distance matrix for BLAST overview tree (Fig. 1); Sheet 4-percent identity of FJG1 hemerythrins found in other methanotrophs; Sheet 5-summary of previous research regarding prokaryotic hemerythrins; Sheet 6 - analyzed proteome data.

**Supp. Movie.** MP4 showing animated overlay of AlphaFold structure for Bhr-00 and crystal structure of Bhr-Bath proteins.

