## supplemental figures for "Diverse bacteriohemerythrin genes of *Methylomonas denitrificans* FJG1 provide insight into the survival and activity of methanotrophs in low oxygen ecosystems"

**Supplemental Figure S1.** Circular alignment of methanotroph genomes created with Blast Ring Image Generator (BRIG). The location of *bhr* genes in *M. denitrificans* FJG1 and other methanotroph genomes are designated by colored bars.

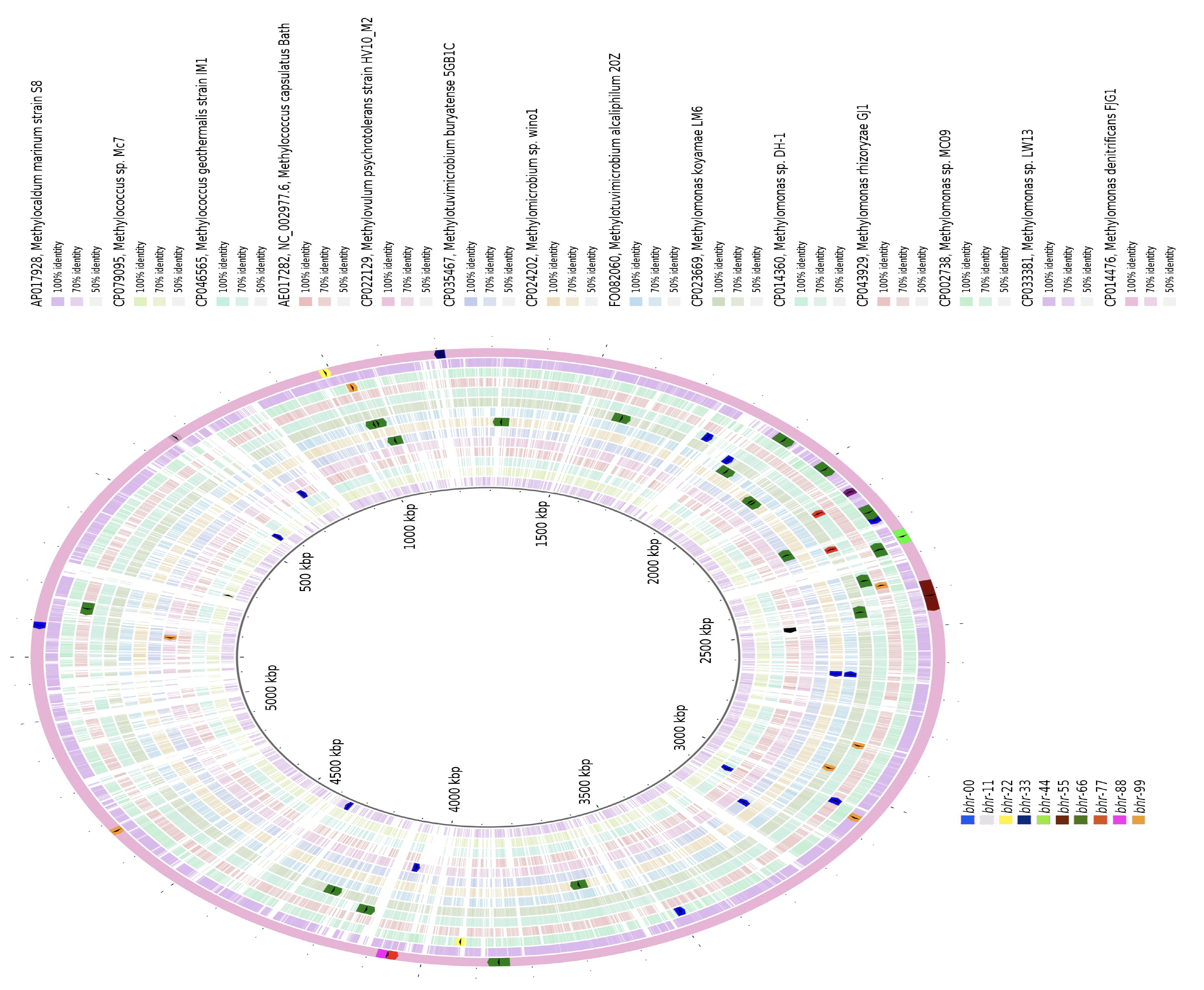

**Supplemental Figure S2.** Alignment of Bhr-00 protein sequence from *Methylomonas denitrificans* FJG1 with Bhr-Bath protein sequence from *Methylococcus capsulatus* Bath.

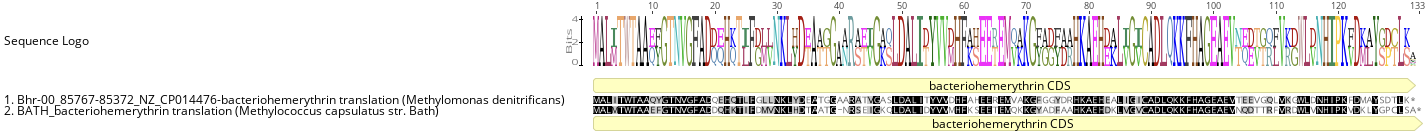

**Supplemental Figure 3**. mRNA and protein expression levels for *bhr-*22, -33, -44, and -55 and gene neighborhoods for cells experiencing high (24 h) and low (48 h) oxygen levels in *M. denitrificans* FJG1 cultures (Supp. Fig. 5). The levels of mRNA expression (in TPM) for NMS (red/orange/yellow heatmap), protein expression (in normalized NSAF) for NMS (pink/purple heatmap) and mRNA expression (in TPM) for AMS (green/yellow heatmap) for *bhr* and neighboring genes are shown for cells extracted at 24 and 48 h time points from left to right. Gene neighbourhood members are shown on the left in their operon structures and are colour-coded by functional group as per the legend. Up arrows indicate genes encoded on the forward DNA strand, and down arrows indicate genes encoded on the reverse DNA strand.

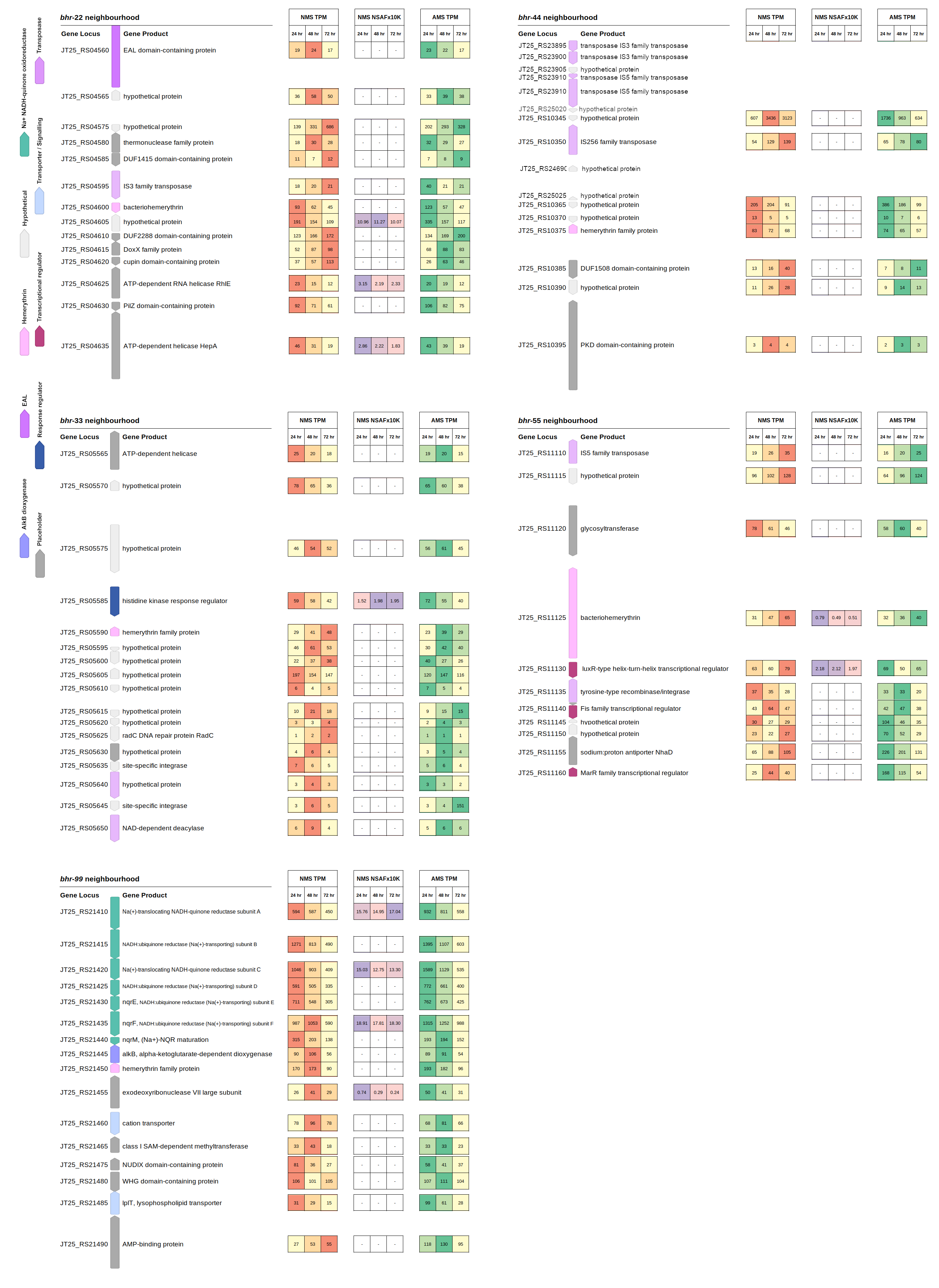

**Supplemental Figure S4.** Gene neighborhoods of *bhr*-66 of non-methanotroph genomes.

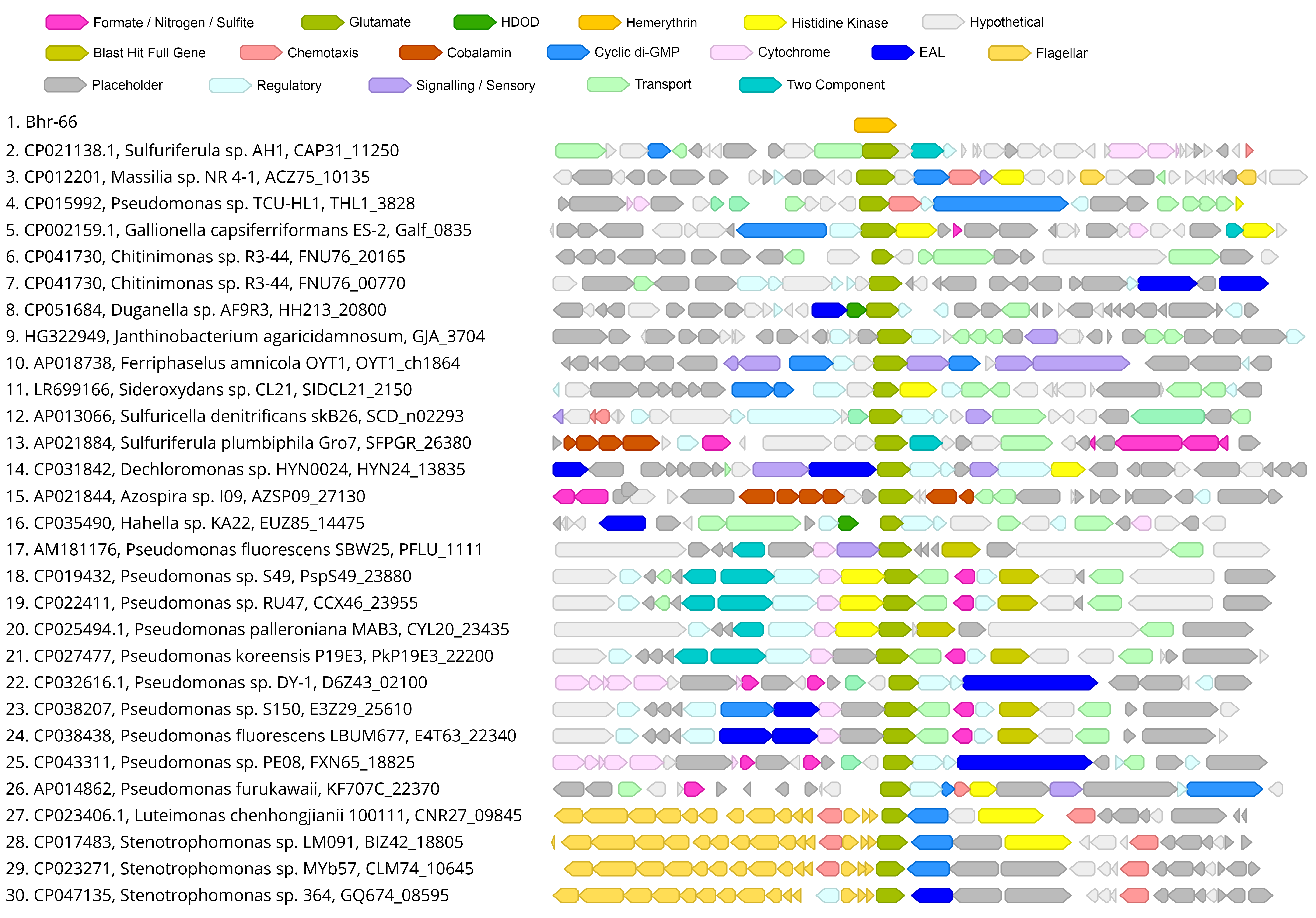

**Supplemental Figure S5.** Growth curve of M. denitrificans FJG1 on NMS or AMS media. (A) Growth over time. (B) Oxygen consumption over time. Reprinted with permission from: Kits KD, Klotz MG, Stein LY. 2015. Methane oxidation coupled to nitrate reduction under hypoxia by the Gammaproteobacterium *Methylomonas denitrificans*, sp. nov. type strain FJG1. Environ Microbiol 17:3219-3232.

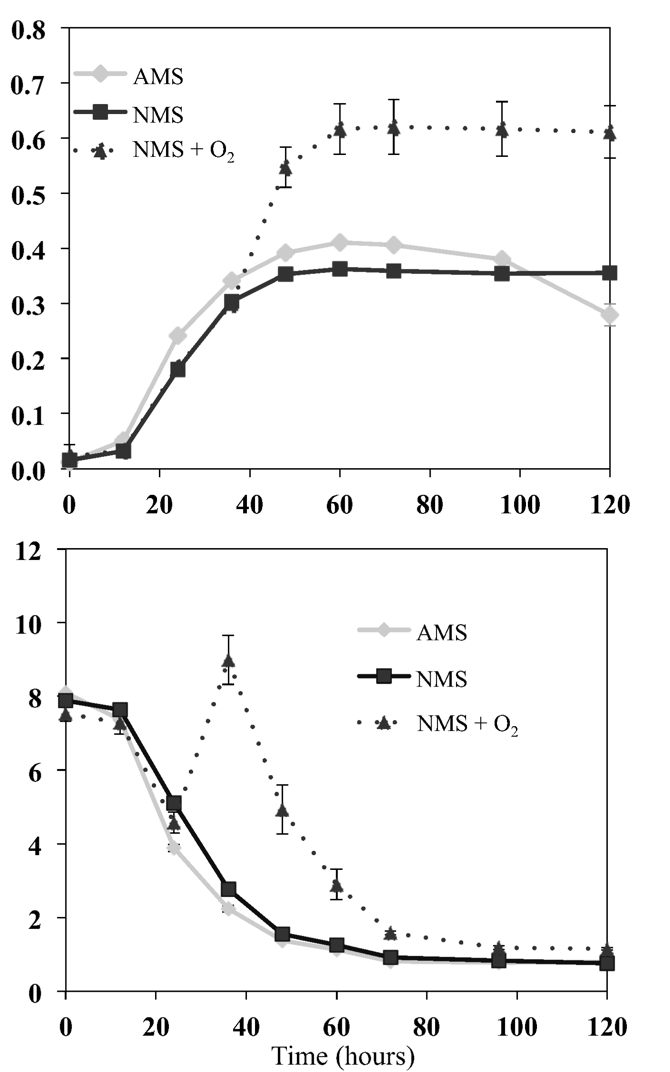

O.D. (600 nm)

O_2_ (mM)

**Supplemental Table 1.** Differential expression of neighboring genes upstream and downstream of the *bhr-*00, -11, -66, and -77/-88 homologues from *M. denitrificans* FJG1 based on RNAseq data at 24 h versus 48 h of growth in NMS and AMS media.

| Gene | Annotation | Locus_tag | size |  |  | Nitrate Treatment (NMS) | | | | |  | Ammonium Treatment (AMS) | | | | |
| --- | --- | --- | --- | --- | --- | --- | --- | --- | --- | --- | --- | --- | --- | --- | --- | --- |
|  |  |  |  |  |  | Raw Expression (TPM) | | Differential Expression 24h vs 48h | | |  | Raw Expression (TPM) | | Differential Expression 24h vs 48h | | |
|  |  |  |  |  |  | 24h | 48h | log2 ratio | adjusted p-value | norm log2 ratio |  | 24h | 48h | log2 ratio | adjusted p-value | norm log2 ratio |
| **bhr-00 biotin neighbourhood** | | | | |  |  |  |  |  |  |  |  |  |  |  |  |
|  | twin transmembrane helix small protein | JT25_RS00355 | 261 | F |  | 79.29 | 34.01 | -1.09 | 0.000 | 0.89 |  | 125.15 | 56.68 | -0.82 | 0.000 | 0.48 |
|  | cytochrome c oxidase subunit 3 | JT25_RS00360 | 876 | R |  | 460.92 | 282.04 | -0.75 | 0.000 | 1.23 |  | 392.14 | 388.57 | 0.16 | 0.000 | 1.46 |
|  | cytochrome c oxidase assembly protein | JT25_RS00365 | 543 | R |  | 1,941.07 | 1,120.97 | -0.97 | 0.000 | 1.01 |  | 1,900.11 | 1,421.06 | -0.33 | 0.000 | 0.98 |
| *ctaD* | cytochrome c oxidase subunit I | JT25_RS00370 | 1617 | R |  | 1,103.57 | 597.90 | -1.01 | 0.000 | 0.97 |  | 875.64 | 771.26 | -0.19 | 0.000 | 1.11 |
| *coxB* | cytochrome c oxidase subunit II | JT25_RS00375 | 1131 | R |  | 1,545.43 | 1,133.14 | -0.60 | 0.000 | 1.37 |  | 1,458.49 | 1,556.46 | 0.07 | 0.000 | 1.37 |
| *bhr-00* | hemerythrin family protein | JT25_RS00380 | 396 | R |  | 4,657.50 | 13,150.82 | 1.35 | 0.000 | 3.33 |  | 7,706.16 | 28,626.86 | 1.91 | 0.000 | 3.21 |
| *bioD* | dethiobiotin synthase | JT25_RS00385 | 696 | R |  | 29.97 | 22.39 | -0.38 | 0.071 | 1.60 |  | 23.62 | 25.97 | -0.10 | 0.517 | 1.20 |
| *bioC* | malonyl-ACP O-methyltransferase BioC | JT25_RS00390 | 789 | R |  | 30.27 | 24.52 | -0.66 | 0.002 | 1.32 |  | 28.87 | 21.38 | -0.29 | 0.041 | 1.01 |
| *bioH* | pimeloyl-ACP methyl ester esterase BioH | JT25_RS00395 | 762 | R |  | 51.20 | 38.11 | -0.66 | 0.000 | 1.32 |  | 76.59 | 36.15 | -0.20 | 0.095 | 1.10 |
| *bioF* | 8-amino-7-oxononanoate synthase | JT25_RS00400 | 1161 | R |  | 34.95 | 33.30 | -0.24 | 0.111 | 1.74 |  | 44.40 | 33.37 | -0.13 | 0.220 | 1.17 |
| *bioB* | biotin synthase BioB | JT25_RS00405 | 990 | R |  | 35.14 | 43.71 | -0.07 | 0.751 | 1.91 |  | 30.27 | 36.99 | 0.01 | 0.910 | 1.31 |
| **bhr-11 nitrogen neighbourhood** | | | | |  |  |  |  |  |  |  |  |  |  |  |  |
| *narA* | nitrate reductase subunit alpha | JT25_RS02985 | 3735 | F |  | 32.81 | 86.13 | 1.03 | 0.000 | 3.00 |  | 35.84 | 19.24 | -0.85 | 0.000 | 0.45 |
| *narH* | nitrate reductase subunit beta | JT25_RS02990 | 1596 | F |  | 42.38 | 105.88 | 0.94 | 0.000 | 2.91 |  | 43.79 | 25.92 | -0.91 | 0.000 | 0.39 |
| *narJ* | nitrate reductase molybdenum  cofactor assembly chaperone | JT25_RS02995 | 720 | F |  | 38.92 | 79.57 | 0.99 | 0.000 | 2.97 |  | 32.73 | 28.02 | -0.49 | 0.000 | 0.81 |
| *narI* | respiratory nitrate reductase subunit gamma | JT25_RS03000 | 678 | F |  | 57.48 | 142.69 | 1.25 | 0.000 | 3.22 |  | 59.90 | 36.01 | -0.99 | 0.000 | 0.31 |
|  | multicopper oxidase  Domain-containing protein | JT25_RS03005 | 1059 | F |  | 26.22 | 69.79 | 1.16 | 0.000 | 3.13 |  | 31.77 | 22.32 | -0.23 | 0.082 | 1.07 |
|  | hypothetical protein | JT25_RS03010 | 219 | F |  | 97.95 | 199.99 | 1.74 | 0.000 | 3.72 |  | 128.30 | 61.83 | -0.63 | 0.000 | 0.67 |
| *bhr-11* | hemerythrin domain-containing protein | JT25_RS03015 | 333 | F |  | 38.22 | 62.58 | 1.17 | 0.000 | 3.15 |  | 75.47 | 23.55 | -1.14 | 0.000 | 0.16 |
|  | HPP family protein | JT25_RS03020 | 1185 | F |  | 45.08 | 54.54 | 0.57 | 0.000 | 2.55 |  | 33.86 | 44.21 | 0.12 | 0.188 | 1.42 |
|  | NAD(P)/FAD-dependent oxidoreductase | JT25_RS03025 | 1230 | R |  | 51.21 | 66.18 | 0.05 | 0.568 | 2.03 |  | 39.95 | 36.87 | 0.43 | 0.000 | 1.73 |
| *phzF* | PhzF family phenazine  Biosynthesis protein | JT25_RS03030 | 921 | R |  | 50.59 | 61.40 | 0.04 | 0.707 | 2.02 |  | 20.63 | 31.09 | 0.31 | 0.013 | 1.61 |
| *nirD* | nitrite reductase small subunit D | JT25_RS03035 | 336 | R |  | 40.85 | 50.84 | 0.41 | 0.008 | 2.39 |  | 18.39 | 28.08 | 0.32 | 0.133 | 1.62 |
| *nirB* | nitrite reductase large subunit B | JT25_RS03040 | 2541 | R |  | 21.28 | 55.98 | 0.75 | 0.000 | 2.72 |  | 14.80 | 11.33 | 0.56 | 0.000 | 1.86 |
|  | formate/nitrite transporter family protein | JT25_RS03045 | 810 | F |  | 22.93 | 130.56 | 2.08 | 0.000 | 4.06 |  | 8.05 | 5.42 | -0.82 | 0.001 | 0.48 |
|  | bifunctional protein-serine/threonine kinase/phosphatase | JT25_RS03050 | 1851 | F |  | 14.22 | 46.66 | 1.34 | 0.000 | 3.32 |  | 5.07 | 5.22 | -0.24 | 0.235 | 1.06 |
| *narK* | NarK family nitrate/nitrite MFS transporter | JT25_RS03055 | 1476 | F |  | 7.48 | 10.10 | 0.91 | 0.000 | 2.89 |  | 1.86 | 2.42 | 0.10 | 0.772 | 1.41 |
| **bhr-66 neighbourhood** | | | | |  |  |  |  |  |  |  |  |  |  |  |  |
| *lexA* | transcriptional repressor LexA | JT25_RS17770 | 618 | R |  | 6.57 | 12.02 | 0.62 | 0.103 | 2.60 |  | 10.07 | 8.74 | -0.49 | 0.053 | 0.81 |
|  | response regulator | JT25_RS17775 | 807 | R |  | 21.14 | 25.57 | 0.01 | 0.965 | 1.99 |  | 24.80 | 22.30 | -0.42 | 0.003 | 0.88 |
|  | DUF447 family protein | JT25_RS17780 | 564 | F |  | 132.49 | 242.09 | 0.71 | 0.000 | 2.68 |  | 155.51 | 222.25 | 0.64 | 0.000 | 1.94 |
|  | hypothetical protein | JT25_RS17785 | 687 | F |  | 100.03 | 139.33 | 0.36 | 0.000 | 2.34 |  | 97.49 | 123.79 | 0.06 | 0.453 | 1.36 |
|  | EAL domain-containing protein | JT25_RS17790 | 3303 | F |  | 52.09 | 49.09 | -0.25 | 0.000 | 1.73 |  | 66.52 | 44.21 | -0.26 | 0.000 | 1.04 |
|  | AEC family transporter | JT25_RS17795 | 924 | F |  | 52.51 | 36.70 | -0.56 | 0.000 | 1.42 |  | 40.93 | 42.50 | -0.22 | 0.021 | 1.08 |
|  | competence/damage-inducible protein A | JT25_RS17800 | 1227 | F |  | 16.21 | 18.39 | 0.09 | 0.694 | 2.07 |  | 22.13 | 19.04 | 0.24 | 0.091 | 1.54 |
|  | ATP-binding cassette domain-containing protein | JT25_RS17805 | 1887 | F |  | 35.74 | 43.64 | 0.03 | 0.822 | 2.01 |  | 53.44 | 44.50 | -0.19 | 0.004 | 1.11 |
|  | hypothetical protein | JT25_RS17810 | 420 | R |  | 68.69 | 73.58 | 0.09 | 0.633 | 2.07 |  | 68.77 | 68.53 | -0.25 | 0.027 | 1.05 |
| *bhr-66* | bacteriohemerythrin | JT25_RS17815 | 1785 | F |  | 48.44 | 63.83 | 0.28 | 0.002 | 2.26 |  | 45.44 | 53.34 | 0.17 | 0.013 | 1.47 |
|  | response regulator | JT25_RS17820 | 1320 | F |  | 39.68 | 45.03 | -0.05 | 0.763 | 1.93 |  | 41.03 | 38.23 | -0.04 | 0.706 | 1.26 |
| *ppk2* | polyphosphate kinase 2 | JT25_RS17825 | 783 | F |  | 36.87 | 50.14 | 0.49 | 0.002 | 2.47 |  | 37.17 | 47.23 | 0.07 | 0.542 | 1.37 |
|  | hypothetical protein | JT25_RS17830 | 645 | R |  | 86.78 | 75.61 | -0.24 | 0.072 | 1.74 |  | 111.16 | 110.81 | 0.23 | 0.003 | 1.53 |
|  | hypothetical protein | JT25_RS17835 | 2244 | R |  | 55.70 | 67.73 | 0.19 | 0.013 | 2.17 |  | 91.41 | 123.89 | 0.50 | 0.000 | 1.80 |
|  | hypothetical protein | JT25_RS17840 | 1140 | R |  | 7.31 | 17.17 | 0.83 | 0.003 | 2.80 |  | 9.25 | 13.59 | 0.27 | 0.113 | 1.57 |
|  | hypothetical protein | JT25_RS17845 | 561 | F |  | 1.96 | 1.29 | -0.23 | 0.782 | 1.75 |  | 1.00 | 1.59 | 0.30 | 0.596 | 1.60 |
| *gspD* | type II secretion system secretin GspD | JT25_RS17850 | 1908 | F |  | 1.38 | 4.89 | 0.83 | 0.230 | 2.81 |  | 0.85 | 1.14 | 0.18 | 0.682 | 1.48 |
| *gspE* | type II secretion system ATPase GspE | JT25_RS17855 | 1512 | F |  | 1.27 | 2.08 | 0.21 | 0.743 | 2.19 |  | 0.95 | 1.57 | 0.35 | 0.409 | 1.66 |
| **bhr-77, bhr-88 chemotaxis neighbourhood** | | | | |  |  |  |  |  |  |  |  |  |  |  |  |
|  | chemotaxis response regulator protein-glutamate methylesterase | JT25_RS18665 | 1095 | R |  | 44.02 | 103.47 | 1.08 | 0.000 | 3.06 |  | 46.36 | 64.11 | 0.54 | 0.000 | 1.84 |
| *cheD* | chemotaxis protein CheD | JT25_RS18670 | 597 | R |  | 34.68 | 163.80 | 1.96 | 0.000 | 3.94 |  | 70.61 | 80.00 | 0.33 | 0.001 | 1.63 |
| cheR | Protein-glutamate O-methyltransferase | JT25_RS18675 | 807 | R |  | 58.08 | 199.36 | 1.55 | 0.000 | 3.53 |  | 53.85 | 101.73 | 0.65 | 0.000 | 1.95 |
|  | hypothetical protein | JT25_RS18680 | 1209 | F |  | 186.80 | 602.19 | 1.59 | 0.000 | 3.57 |  | 196.48 | 406.79 | 0.92 | 0.000 | 2.22 |
|  | response regulator | JT25_RS18685 | 369 | F |  | 312.41 | 1,079.75 | 1.64 | 0.000 | 3.62 |  | 417.07 | 749.70 | 1.04 | 0.000 | 2.34 |
|  | STAS domain-containing protein | JT25_RS18690 | 333 | F |  | 214.74 | 532.60 | 1.21 | 0.000 | 3.19 |  | 238.96 | 386.06 | 0.87 | 0.000 | 2.17 |
| *cheA* | chemotaxis protein CheA | JT25_RS18695 | 2202 | F |  | 127.43 | 297.19 | 1.08 | 0.000 | 3.06 |  | 111.08 | 191.32 | 0.52 | 0.000 | 1.82 |
|  | HAMP domain-containing protein | JT25_RS18700 | 2034 | F |  | 176.46 | 453.17 | 1.21 | 0.000 | 3.19 |  | 200.61 | 324.35 | 0.71 | 0.000 | 2.02 |
| *cheW* | purine-binding chemotaxis protein CheW | JT25_RS18705 | 546 | F |  | 209.45 | 446.22 | 0.99 | 0.000 | 2.97 |  | 297.34 | 336.69 | 0.41 | 0.000 | 1.71 |
| *bhr-77* | bacteriohemerythrin | JT25_RS18710 | 405 | R |  | 107.16 | 176.26 | 0.62 | 0.000 | 2.60 |  | 146.80 | 119.38 | 0.07 | 0.470 | 1.37 |
| *bhr-88* | hemerythrin family protein | JT25_RS18715 | 414 | R |  | 50.66 | 80.73 | 0.59 | 0.000 | 2.57 |  | 64.90 | 73.10 | 0.22 | 0.071 | 1.52 |
|  | methyl-accepting chemotaxis protein | JT25_RS18720 | 1878 | R |  | 21.25 | 39.93 | 0.51 | 0.001 | 2.49 |  | 24.06 | 33.64 | 0.23 | 0.007 | 1.53 |
|  | formate--tetrahydrofolate ligase | JT25_RS18725 | 1674 | R |  | 61.58 | 69.40 | 0.13 | 0.181 | 2.11 |  | 82.37 | 74.66 | -0.29 | 0.000 | 1.01 |
| **Normalization** | | | | |  |  |  |  |  |  |  |  |  |  |  |  |
| *rpoB* | DNA-directed RNA polymerase  Subunit beta | JT25_RS14695 | 4077 | F |  | 418.02 | 113.34 | -1.98 | 0.000 |  |  | 421.34 | 162.99 | -1.30 | 0.000 |  |
